# Rapid Ultrastructural Changes of PSD and Extrasynaptic Axon-spine Interface Membrane during LTP Induced in Single Dendritic Spine

**DOI:** 10.1101/840629

**Authors:** Ye Sun, Michael Smirnov, Naomi Kamasawa, Ryohei Yasuda

## Abstract

Structural plasticity of dendritic spines is considered to be the basis of synaptic plasticity, learning and memory. Here, we performed ultrastructural analysis of spines undergoing LTP using a novel high throughput correlative light-electron microscopy approach. We found that the PSD displays rapid (< 3 min) reorganization of its nanostructure, including perforation and segmentation. This increased structural complexity is maintained over intermediate and late phases of LTP (20 and 120 min). In a few spines, segmented PSDs are connected to different presynaptic terminals, producing a multi-innervated spine in the intermediate and late phases. In addition, the area of extrasynaptic axon-spine interface (eASI) displayed a pronounced, rapid and sustained increase. Finally, presynaptic vesicle number increased slowly and became significantly higher at late phases of LTP. These rapid ultrastructural changes in PSD and surrounding membrane, together with the slow increase in presynaptic vesicle number, likely support the rapid and sustained increase in synaptic transmission during LTP.

## Introduction

Dendritic spines are the major sites for receiving excitatory synaptic inputs and play important roles in neuronal signal transduction, memory storage and neuronal circuit organization. Plasticity of spine structure is considered to play critical roles in functional plasticity of synaptic transmission, learning and memory (Sala and Segal, 2014). In particular, it has been reported that activity-dependent spine enlargement is correlated with LTP (Kopec et al., 2006; Lang et al., 2004; Matsuzaki et al., 2004) and memory formation (Hayashi-Takagi et al., 2015). Ultrastructural correlates of LTP have been also studied extensively using electron micrographs of LTP-induced brain slices under a condition where a large fraction of spines are electrically stimulated. These studies revealed that LTP is correlated with, on average, an enlargement in spine and PSD size, a higher number of concave spine profiles, an increase in perforated PSD, and an increase in polyribosome containing spines (Desmond and Levy, 1986; Van Harreveld and Fifkova, 1975; Harris et al., 1992; Ostroff et al., 2002). Furthermore, LTP has been associated with generation of multi-innervating spines and boutons (Giese et al., 2015; Toni et al., 1999).

One difficulty of ultrastructural studies of spine structural plasticity is to identify which spines have undergone synaptic plasticity. To address this, single spines with LTP induction using 2-photon (2P) glutamate uncaging have been visualized under transmission electron microscopy (TEM) after serial ultra-thin sectioning. This allows correlation between light and electron microscopy and opens a possibility to perform ultrastructural analyses on spines undergoing plasticity (Bosch et al., 2014; Meyer et al., 2014). It has been found that, while spine volume increase occurs rapidly, PSD area and bouton volume increase slowly over several hours (Bosch et al., 2014; Meyer et al., 2014). This apparently contradicts with a rapid increase of synaptic transmission during LTP (Bosch et al., 2014; Matsuzaki et al., 2004). However, the throughput of these methods has been low, and thus detailed analysis of PSD morphology and presynaptic vesicles have not been done.

In addition to these studies, PSD structure and plasticity has been extensively studied using optical super-resolution microscopy (Hruska et al., 2018; MacGillavry et al., 2013; Masch et al., 2018; Nair et al., 2013; Wegner et al., 2018). These studies suggest that, in dissociated culture, PSD is made of isolated nanoclusters (Hruska et al., 2018; MacGillavry et al., 2013; Nair et al., 2013), which increases in number during chemically-induced LTP (Hruska et al., 2018; MacGillavry et al., 2013). However, a recent in vivo study using stimulated emission depletion (STED) microscopy suggested the PSD has a simple, round/ovoid structure, or complex but continuous shape, but does not form separate clusters (Masch et al., 2018). Thus, overall, how the PSD, surrounding membrane and presynaptic sites change their nanometer-level structure during LTP still remain elusive.

Here, we demonstrate that single spines stimulated and imaged under a 2P microscope can be efficiently re-imaged with automated tape-collecting an ultramicrotome (ATUMtome) sectioning and scanning EM (SEM). This technique allows for clear visualization of sub-spine ultrastructures including the PSD, synaptic membrane, and synaptic vesicles (Kamasawa et al., 2015). 3D reconstruction of the stimulated spines showed that the PSD undergoes rapid and sustained structural changes, including perforation and segmentation, while its overall size increases slowly. Occasionally, segmented PSDs are connected to different presynaptic terminals, producing a multi-innervated spine over 20 – 120 min. Associated with changes in the PSD structure, we also observed a pronounced expansion of the extrasynaptic axon-spine interface (eASI), the postsynaptic membrane without an attached PSD but having a typical synaptic cleft (< 40 nm) against presynaptic membrane. This expansion occurs rapidly after LTP induction (2-3 min) and is maintained over a long time (20 min and 120 min). Furthermore, the number of presynaptic vesicles slowly increased and became significantly higher at 120 min. The expansion of eASI, supported by PSD with high structural complexity, could serve as the physical location for transient accumulation of AMPA receptors to support a rapid increase in synaptic transmission in the early phase of LTP, while the slow increase in presynaptic function and PSD size may support late phase LTP.

## Results

### Correlating dendritic spines imaged by 2P microscopy and SEM

We established a high-throughput pipeline to analyze the 3D ultrastructure of spines undergoing LTP in organotypic hippocampal slices. Spine structural plasticity was first imaged using 2p microscopy, and then identified in EM by combining pre-embedding immuno-labeling of GFP with ATUMtome serial sectioning and SEM imaging (Fig. 1A-E; whole flow in S1). We induced LTP at single spines with glutamate uncaging by delivering 2P laser pulses located ~0.5 µm from the tip of the spine head in EGFP-expressing CA pyramidal neurons in organotypic hippocampal slices (Fig.1A-C, S1A). These slices were then fixed at different time points after uncaging stimulation to capture the structural changes at different LTP phases: early phase (2-3 min), intermediate phase (20 min) and late phase (120 min) (Fig. S1B). After immuno-staining of EGFP with nanogold particles, we performed silver enhancement, heavy metal staining, dehydration and resin embedding. Using ATUMtome, we cut and collect ultrathin sections (50 nm), which were imaged in SEM at low resolution (2 µm/pixel) and intermediate resolution (30 nm/pixel) (Fig. S1C, D1-D2). Following identification of target neurons and secondary dendrites with the help of nanogold labeling, we imaged spines that underwent LTP (LTP spine) and spines located within 20 µm of the stimulated spine on the same dendrite (control spine) with high resolution (4 nm/pixel) (Fig. 1D, S1D3). 3D morphology of dendritic spines and PSDs were reconstructed manually from the sections and subjected to further analyses (Fig. 1E, S1D4).

**Figure 1.**
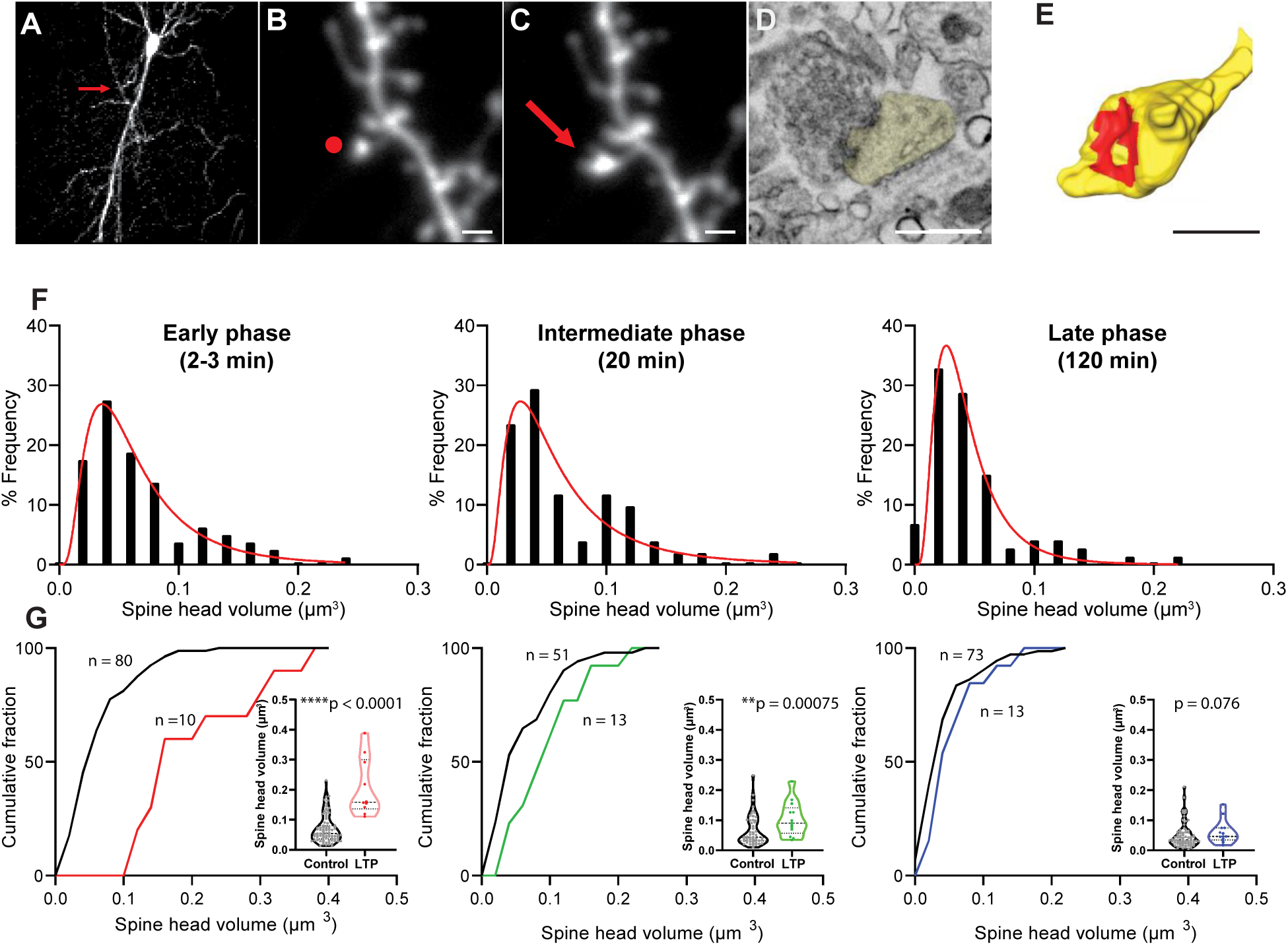
LTP induces spine expansion. **(A)** 2P image of a GFP positive CA1 pyramidal neuron in organotypic hippocampal slice selected for glutamate uncaging. Red arrow points to the location of target spine. Scale bar: 20 µm. **(B-C)** 2P images of a dendritic spine before **(B)** and after **(C)** 2P glutamate uncaging stimulation. Red dot in **(B)** shows the uncaging position, and red arrow in **(C)** points to the enlarged dendritic spine. Scale bar: 1 µm. **(D)** 4 nm/pixel high resolution image taken with InlensDuo detector shows details of modulated dendritic spine ultrastructure. Spine head colored in yellow. Scale bar: 0.5 µm. **(E)** 3D reconstruction of the target dendritic spine. Spine head colored in yellow, and PSD colored in red. Scale bar: 0.5 µm. **(F)** Distribution of raw spine head volume for control spines in early, intermediate and late phases of LTP. **(G)** Cumulative distribution of raw spine head volume for control and LTP spines in early, intermediate and late phases of LTP. Welch’s t-test on log-transformed data.

The volume of control spines was 0.061 ± 0.003 µm^3^ (mean ± sem, n = 204), consistent with previous studies (Harris et al., 1992). From fluorescence changes following LTP and the volume of LTP spines in EM, we estimated that the average volume of LTP spines before LTP (0.063 ± 0.006 µm^3^; mean ± sem, n = 36) was similar to the volume of control spines, suggesting that our LTP spines represent average spines in terms of their volume before LTP. We found that control spine volume distribution is skewed from normal distribution and can be expressed by log-normal function (Fig. 1F) (Loewenstein et al., 2011). Thus for the rest of the paper, we performed parametric statistics after log-transformation for datasets which follow a log-normal distribution. In addition, since ultrastructure of spines may slowly change over time in ACSF over hours, we compared only between LTP spines and surrounding spines, fixed at the same time point. As expected, we found that LTP spines showed significantly larger size during the early phase and immediate phase compared with control spines. However, due to the large variability in spine volume, the spine size during the late phase was not statistically larger, although they showed enlargement in fluorescent microscopy (Fig. 1G, Table 1). Similarly, we found that spine surface area increased after LTP during early and immediate, as well as in the late phase (Table 1).

**Table 1.**
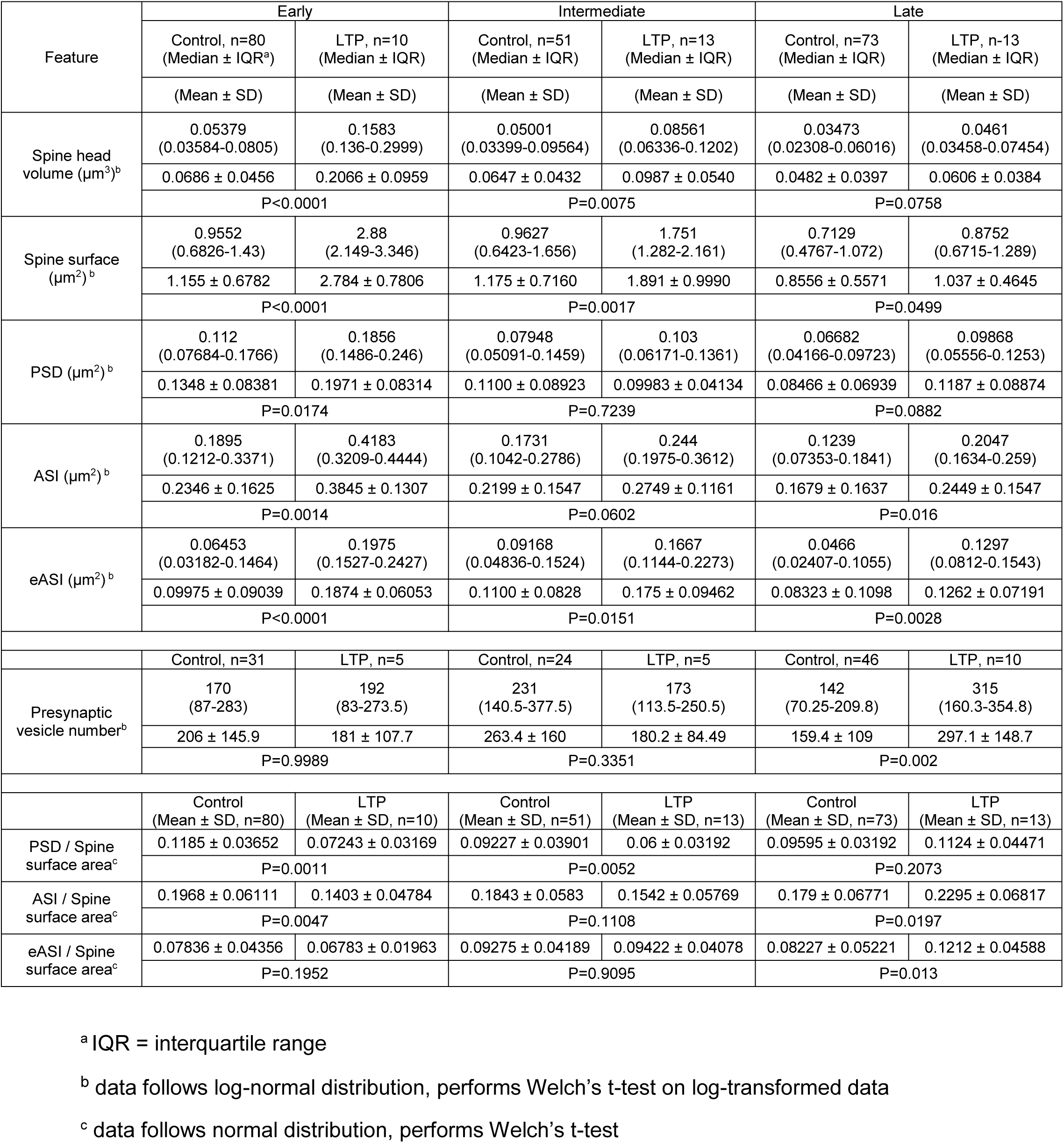
Comparison of the ultrastructures of dendritic spines in control and LTP conditions at early, intermediate and late phases.

| Feature | Early |  | Intermediate |  | Late |  |
| --- | --- | --- | --- | --- | --- | --- |
|  | Control, n=80<br>(Median ± IQR<sup>a</sup>) | LTP, n=10<br>(Median ± IQR) | Control, n=51<br>(Median ± IQR) | LTP, n=13<br>(Median ± IQR) | Control, n=73<br>(Median ± IQR) | LTP, n=13<br>(Median ± IQR) |
|  | (Mean ± SD) | (Mean ± SD) | (Mean ± SD) | (Mean ± SD) | (Mean ± SD) | (Mean ± SD) |
| Spine head<br>volume (μm<sup>3</sup>)<sup>b</sup> | 0.05379<br>(0.03584-0.0805) | 0.1583<br>(0.136-0.2999) | 0.05001<br>(0.03399-0.09564) | 0.08561<br>(0.06336-0.1202) | 0.03473<br>(0.02308-0.06016) | 0.0461<br>(0.03458-0.07454) |
|  | 0.0686 ± 0.0456 | 0.2066 ± 0.0959 | 0.0647 ± 0.0432 | 0.0987 ± 0.0540 | 0.0482 ± 0.0397 | 0.0606 ± 0.0384 |
|  | P<0.0001 |  | P=0.0075 |  | P=0.0758 |  |
| Spine surface<br>(μm<sup>2</sup>)<sup>b</sup> | 0.9552<br>(0.6826-1.43) | 2.88<br>(2.149-3.346) | 0.9627<br>(0.6423-1.656) | 1.751<br>(1.282-2.161) | 0.7129<br>(0.4767-1.072) | 0.8752<br>(0.6715-1.289) |
|  | 1.155 ± 0.6782 | 2.784 ± 0.7806 | 1.175 ± 0.7160 | 1.891 ± 0.9990 | 0.8556 ± 0.5571 | 1.037 ± 0.4645 |
|  | P<0.0001 |  | P=0.0017 |  | P=0.0499 |  |
| PSD (μm<sup>2</sup>)<sup>b</sup> | 0.112<br>(0.07684-0.1766) | 0.1856<br>(0.1486-0.246) | 0.07948<br>(0.05091-0.1459) | 0.103<br>(0.06171-0.1361) | 0.06682<br>(0.04166-0.09723) | 0.09868<br>(0.05556-0.1253) |
|  | 0.1348 ± 0.08381 | 0.1971 ± 0.08314 | 0.1100 ± 0.08923 | 0.09983 ± 0.04134 | 0.08466 ± 0.06939 | 0.1187 ± 0.08874 |
|  | P=0.0174 |  | P=0.7239 |  | P=0.0882 |  |
| ASI (μm<sup>2</sup>)<sup>b</sup> | 0.1895<br>(0.1212-0.3371) | 0.4183<br>(0.3209-0.4444) | 0.1731<br>(0.1042-0.2786) | 0.244<br>(0.1975-0.3612) | 0.1239<br>(0.07353-0.1841) | 0.2047<br>(0.1634-0.259) |
|  | 0.2346 ± 0.1625 | 0.3845 ± 0.1307 | 0.2199 ± 0.1547 | 0.2749 ± 0.1161 | 0.1679 ± 0.1637 | 0.2449 ± 0.1547 |
|  | P=0.0014 |  | P=0.0602 |  | P=0.016 |  |
| eASI (μm<sup>2</sup>)<sup>b</sup> | 0.06453<br>(0.03182-0.1464) | 0.1975<br>(0.1527-0.2427) | 0.09168<br>(0.04836-0.1524) | 0.1667<br>(0.1144-0.2273) | 0.0466<br>(0.02407-0.1055) | 0.1297<br>(0.0812-0.1543) |
|  | 0.09975 ± 0.09039 | 0.1874 ± 0.06053 | 0.1100 ± 0.0828 | 0.175 ± 0.09462 | 0.08323 ± 0.1098 | 0.1262 ± 0.07191 |
|  | P<0.0001 |  | P=0.0151 |  | P=0.0028 |  |
|  | Control, n=31 | LTP, n=5 | Control, n=24 | LTP, n=5 | Control, n=46 | LTP, n=10 |
| Presynaptic<br>vesicle number<sup>b</sup> | 170<br>(87-283) | 192<br>(83-273.5) | 231<br>(140.5-377.5) | 173<br>(113.5-250.5) | 142<br>(70.25-209.8) | 315<br>(160.3-354.8) |
|  | 206 ± 145.9 | 181 ± 107.7 | 263.4 ± 160 | 180.2 ± 84.49 | 159.4 ± 109 | 297.1 ± 148.7 |
|  | P=0.9989 |  | P=0.3351 |  | P=0.002 |  |
|  | Control<br>(Mean ± SD, n=80) | LTP<br>(Mean ± SD, n=10) | Control<br>(Mean ± SD, n=51) | LTP<br>(Mean ± SD, n=13) | Control<br>(Mean ± SD, n=73) | LTP<br>(Mean ± SD, n=13) |
| PSD / Spine<br>surface area<sup>c</sup> | 0.1185 ± 0.03652 | 0.07243 ± 0.03169 | 0.09227 ± 0.03901 | 0.06 ± 0.03192 | 0.09595 ± 0.03192 | 0.1124 ± 0.04471 |
|  | P=0.0011 |  | P=0.0052 |  | P=0.2073 |  |
| ASI / Spine<br>surface area<sup>c</sup> | 0.1968 ± 0.06111 | 0.1403 ± 0.04784 | 0.1843 ± 0.0583 | 0.1542 ± 0.05769 | 0.179 ± 0.06771 | 0.2295 ± 0.06817 |
|  | P=0.0047 |  | P=0.1108 |  | P=0.0197 |  |
| eASI / Spine<br>surface area<sup>c</sup> | 0.07836 ± 0.04356 | 0.06783 ± 0.01963 | 0.09275 ± 0.04189 | 0.09422 ± 0.04078 | 0.08227 ± 0.05221 | 0.1212 ± 0.04588 |
|  | P=0.1952 |  | P=0.9095 |  | P=0.013 |  |

<sup>a</sup> IQR = interquartile range
<sup>b</sup> data follows log-normal distribution, performs Welch's t-test on log-transformed data
<sup>c</sup> data follows normal distribution, performs Welch's t-test

### PSD reorganization after LTP induction

We next reconstructed the PSD in 3D for each spine (Fig. 2A, B). The PSD area size in LTP spines were significantly larger than that in control spines in the early time point (Fig. 2C), albeit to a less degree compared with the spine volume and surface area change. The area occupancy of PSD, the fraction of PSD area over the total spine surface area, was reduced in the early and intermediate phases, but becomes similar to control PSD at the late phase of LTP (Fig. 2D). This suggest an increase in PSD area size takes longer time than that in the total spine surface.

**Figure 2.**
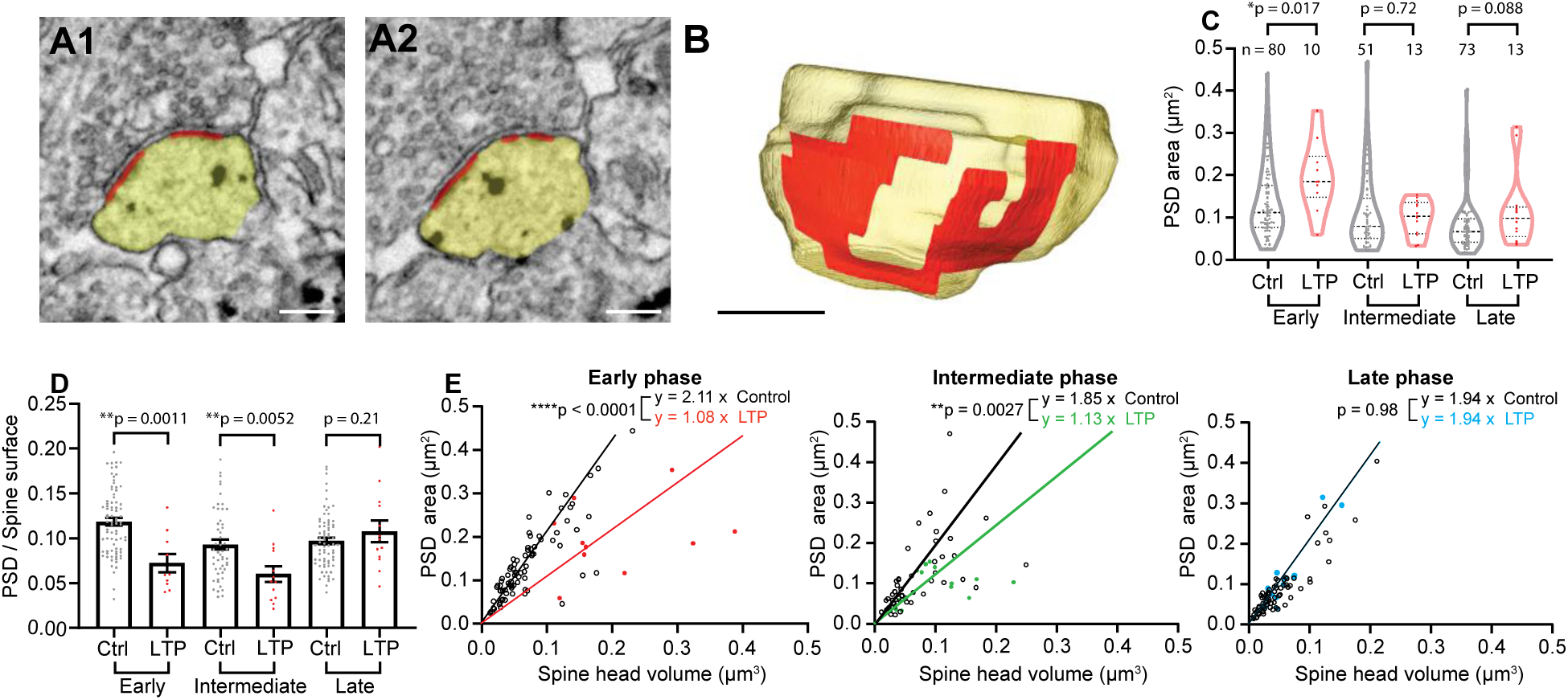
LTP induction disrupts correlation between PSD and spine head volume. **(A-B)** Serial images of an example spine **(A)** and 3D reconstruction of the example spine **(B)** with spine head colored in yellow, and PSD colored in red. Scale bar: 200 nm. **(C)** PSD size at early, intermediate and late phases. Welch’s t-test on log-transformed data. **(D)** Ratio of PSD size to spine surface area at early, intermediate and late phases. Welch’s t-test, mean ± SEM. **(E)** Correlation of PSD area size and spine head volume for control and LTP samples at early, intermediate and late phases of LTP. The regression lines represent an average of PSD area / spine volume in each group. P values are calculated by Welch’s t-test on PSD area / spine volume.

We also analyzed the relationship between spine volume and PSD area (Fig. 2E). In control spines, PSD volume was proportional to the spine size, consistent with previous studies (Bosch et al., 2014; Harris and Stevens, 1989). The slope of the PSD-volume relationship dramatically decreased at the early and intermediate phases, suggesting that the PSD growth was much smaller than the volume changes during these phases. The slope became similar to control at the late phase, consistent with the slow growth of PSD area compared with spine volume increase during LTP (Bosch et al., 2014).

We further analyzed detailed nanometer-scale structure of the PSD (Fig. 3A-H). We observed PSD shapes with previously reported features, including simple, fenestration, horseshoe, and segmentation (Fig. 3A-D). In addition, we also observed spines with irregular shapes (Fig. 3E, F), and with shapes including multiple features (Fig. 3G, H). At all time points, a higher percentage of LTP spines contained PSDs with complex structural features, including fenestration, horseshoe-shape, and segmentation, compared to control spines (Fig. 3I). In the early phase, nearly half of the stimulated spines contained segmented PSD, composed of either 2 or 3 fragments (Fig. 3J). To quantitatively evaluate the complexity of PSD shape, we calculated a complexity index, defined as the ratio of the square of the perimeter and the area, normalized to that of the circular disk shape (perimeter^2^ / (4π area)). The complexity index was significantly higher in LTP spines than control spines from early LTP phase (Fig. 3K). Taken together, our results indicate that a dramatic PSD shape transformation happened immediately after LTP induction. We also observed that, at the intermediate and late time points, some of the segmented PSDs in LTP spines are connected with different presynaptic boutons, forming multi-innervating spines (1 in 13 for both phases), while control spines are nearly exclusively connected with a single presynaptic bouton (0 in 51 for intermediate phase, 2 in 73 for late phase) (Fig. 3L-N).

**Figure 3.**
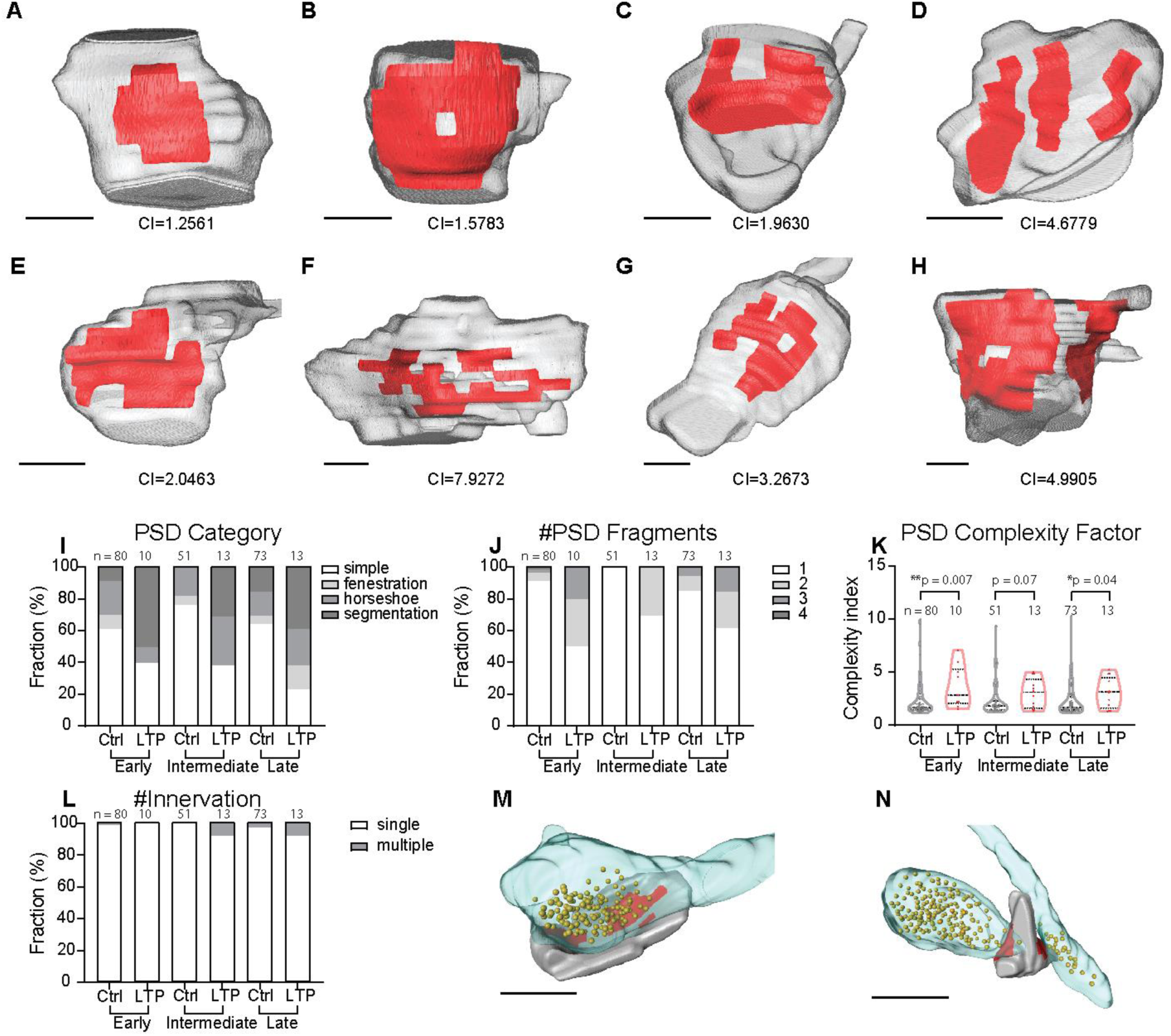
LTP induces increased PSD complexity. **(A-H)** 3D reconstruction of example spines(grey) containing PSD (red) with different complexity indexes (CI) and different shape categories (**(A)** simple, **(B)** fenestration, **(C)** horseshoe, **(D)** segmentation, **(E, F)** irregular horseshoe, **(G,H)** combined shape categories). Scale bar: 200 nm. **(I)** Percentage of spines contains PSD in simple, fenestrated, horseshoe, segmented categories. **(J)** Percentage of spines containing 1-4 PSD segments. **(K)** PSD complexity index at early, intermediate and late phases. Mann-Whitney t-test on log-transformed data. **(L)** Percentage of single- and multi-innervated spines. **(M-N)** 3D reconstruction of example single-innervated spine **(M)** and multi-innervated spine **(N)** with opposing presynaptic boutons and synaptic vesicles. Spines colored in grey, PSD colored in red, presynaptic bouton colored in transparent blue, synaptic vesicles shown as golden balls. Scale bar: 500 nm.

### Extrasynaptic axon-spine interface expansion after LTP induction

Synaptic area is usually defined as the postsynaptic membrane containing the PSD. However, extrasynaptic axon-spine interface (eASI), defined as postsynaptic membrane without an attached PSD but having a typical synaptic cleft (< 40 nm) against presynaptic membrane (Fig. 4A, B) may also be able to support synaptic transmission. We found that eASI expanded immediately after LTP induction, and was maintained at a larger size until the late phase (Fig. 4C). The eASI area occupancy, the fraction of eASI area over the total spine surface area, in LTP spines was similar to control spines in the early and intermediate phases (Fig. 4D). Thus, unlike PSD area, the eASI area size increases immediately with the same rate as the spine surface membrane. Interestingly, the eASI area occupancy became larger than control in the late phase, suggesting that the growth of eASI exceeds the growth of the spine surface area in the late phase.

**Figure 4.**
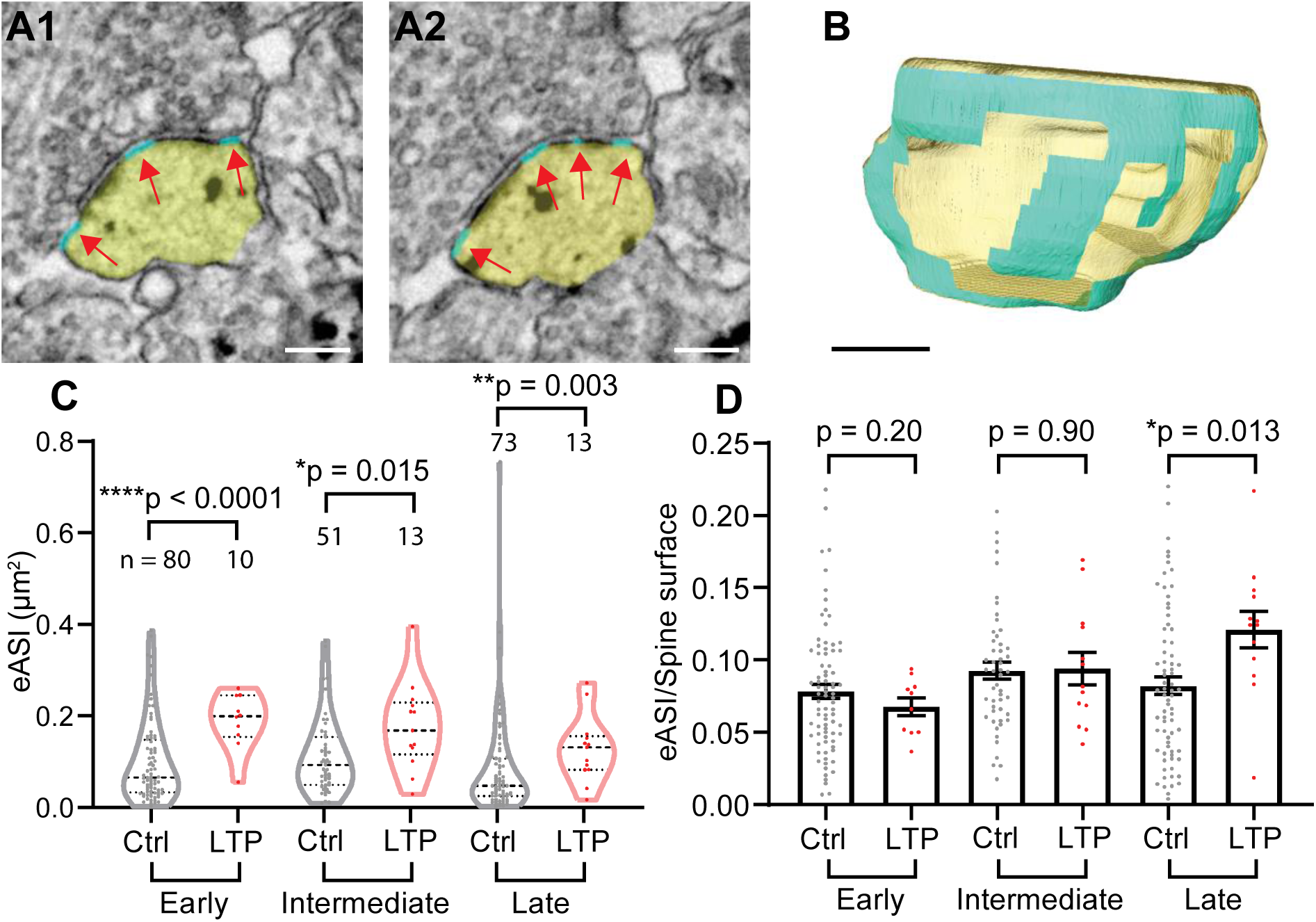
LTP induces sustained expansion of extrasynaptic axon-spine interface (eASI). **(A-B)** Serial images of an example spine **(A)** and 3D reconstruction of the example spine **(B)** with spine head colored in yellow, and eASI area colored in cyan, pointed by red arrows. Scale bar: 200 nm. **(C)** eASI size at early, intermediate and late phases. Welch’s t-test on log-transformed data. **(D)** Ratio of eASI area to spine surface area at early, intermediate, and late phases. Welch’s t-test.

Taken together with PSD area growth, we also calculated the size of total axon-spine interface (ASI = PSD + eASI) (Fig. S2A, B). A rapid and maintained expansion of ASI was observed (Fig. S2C). The area occupancy of ASI (ASI area / spine surface) suggests that, while the degree of expansion is smaller than that of spine surface area in the early phase, it becomes larger in the late phase (Fig. S2D).

### Presynaptic vesicles increase at late LTP phase

We next analyzed the number of presynaptic vesicles as a structural correlate of presynaptic function. The number of presynaptic vesicles in LTP spines was similar to control spines in early and intermediate phases, but higher in the late phase of LTP (Fig. 5), in agreement with literature suggesting presynaptic terminal strength increases more slowly than spine volume change (Meyer et al., 2014).

**Figure 5.**
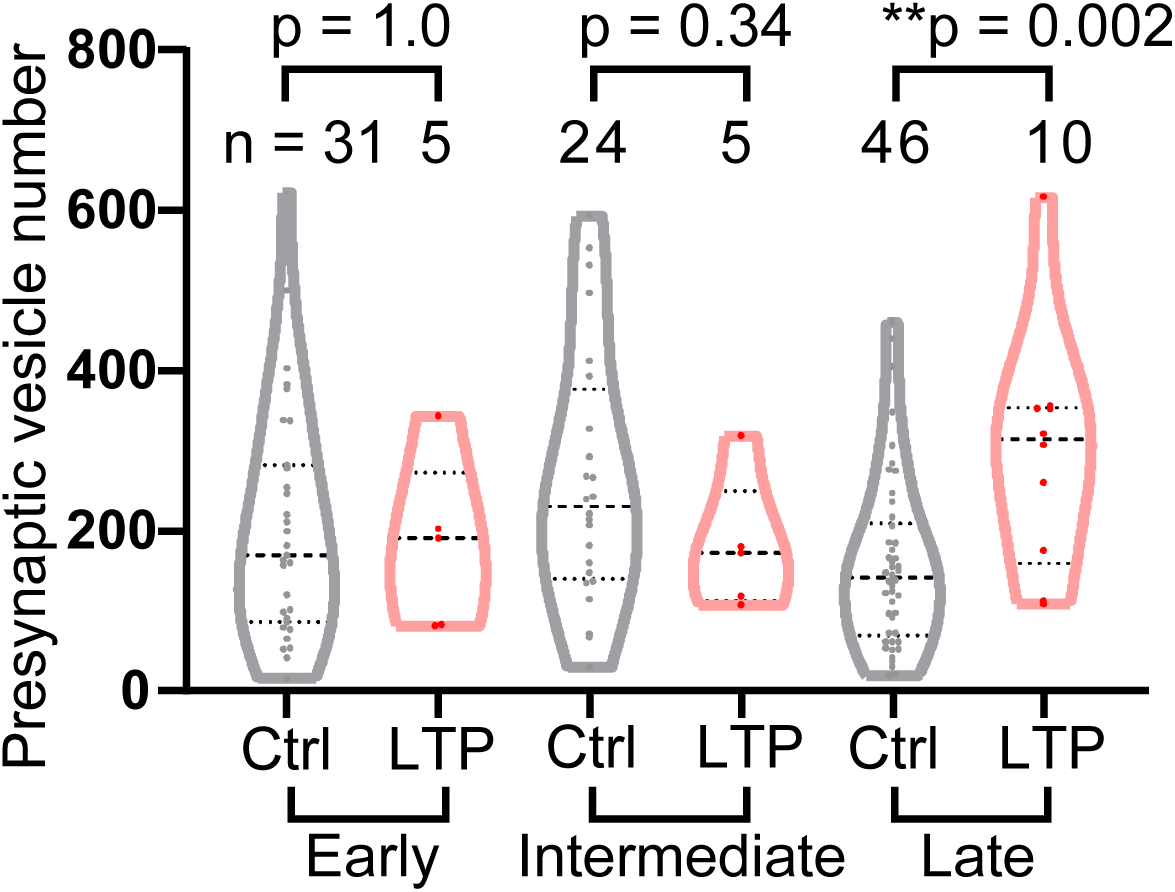
Presynaptic vesicle number increase at late phase LTP. Number of presynaptic vesicles at early, intermediate and late phases. Welch’s t-test on log-transformed data. N is smaller than other parameters, because only a fraction of samples showed clear presynaptic vesicles.

## Discussion

In this study, we developed a high throughput method for analyzing ultrastructural changes of dendritic spines after LTP. Using this method, we found that PSD complexity increases rapidly within 2-3 min by nanometer-scale structural changes including perforation and segmentation. In particular, a large fraction of PSDs showed rapid fragmentation into 2-3 nanoclusters, and a few of them appear to be innervated by multiple boutons. Associated with these changes, PSD size increases during the early phase of LTP, although it is not as pronounced as changes in spine surface area. On the other hand, eASI membrane shows pronounced and sustained expansion, which begins with a similar rate as spine surface area expansion but exceeds the spine surface area growth rate at 120 min. The number of presynaptic vesicles, which is perhaps correlated with presynaptic function, shows a slow increase and became significant 120 min after LTP induction.

Previously ultrastructural analyses following LTP induced by uncaging have also been performed using serial sectioned TEM (Bosch et al., 2014; Meyer et al., 2014). However, perhaps due to low yield, these studies restricted their analyses on PSD area size, spine volume and presynaptic bouton sizes. Our methods using ATUMtome serial sectioning and SEM imaging allowed us to image with relatively high throughput and to perform detailed analyses of nanostructure of PSDs and spines as well as presynaptic vesicle number in synapses undergoing LTP.

It is well known that LTP induction, either by glutamate uncaging or electrical stimulation, is associated with a rapid expansion of spine head volume coupled with an increase in postsynaptic sensitivity (Bosch et al., 2014; Harvey et al., 2008; Lang et al., 2004; Lee et al., 2009; Matsuzaki et al., 2004; Yang et al., 2008a) as well as presynaptic bouton volume change (Meyer et al., 2014). The changes of spine head volume is similar to the onset and time course of electrophysiological LTP indicated by AMPAR-mediated current (Kopec et al., 2006; Matsuzaki et al., 2004). However, it has been reported that potentiation of PSD size is much slower than that of volume change, apparently inconsistent with rapid potentiation of AMPAR-mediated current (Bosch et al., 2014).

Our finding of the rapid complication of PSD nanostructure, associated with expansion of eASI area may be a mechanism underlying rapid recruitment of AMPAR to the PSD. Although eASI membrane does not contain scaffolding proteins to anchor glutamate receptors for regular synaptic transmission, AMPARs are known to exist in extrasynaptic membrane near the PSD (Nair et al., 2013; Tao-Cheng et al., 2011) and likely be able to support synaptic transmission (Franks et al., 2003; Tardin et al., 2003). In particular, extrasynaptic AMPAR may play even larger roles in LTP, since LTP is known to be associated with greater amount of glutamate release (Bekkers and Stevens, 1990; Dolphin et al., 1982; Malinow and Tsien, 1990; Zakharenko et al., 2001) and potentially larger glutamate spillover due to astrocyte withdraw (Henneberger et al., 2019). This idea is consistent with previous electrophysiological experiments showing rapid recruitment of AMPARs to perisynaptic sites before the full expression of LTP (Yang et al., 2008b).

Previous studies suggest that metabotropic receptors (mGluRs) exist in perisynaptic areas within ~60 nm of synapses (Luján et al., 1996). Our structurally defined eASI area thus should include mGluRs. Although mGluRs have been shown to be involved in some forms of LTP (Bashir et al., 1993), blockade of mGluRs does not affect induction of sLTP in our conditions (Matsuzaki et al., 2004; Zhai et al., 2013). However, rapid and pronounced expansion of eASI area observed in our study may cause enhanced mGluR function after LTP induction.

We observed a few spines containing segmented PSDs are innervated by multiple presynaptic boutons. An increase of multi-innervated spines has been reported to be associated with LTP (Giese et al., 2015; Nikonenko et al., 2003; Radwanska et al., 2011). However, probably due to the rare existence of multi-innervated spines (~1% of total spines), it is seldom reported in previous population EM studies that analyzed the ultrastructural correlates for LTP. Since LTP is known to be correlated with larger number of synapses in CA1 neurons (Tominaga-Yoshino et al., 2008; Watson et al., 2016), the rapid multi-innervated spines formation may serve as the basis for new synapse formation during LTP.

Our LTP induction protocol bypasses presynaptic terminal stimulation. However, presynaptic vesicle number still increased, consistent with a previous study showing presynaptic bouton size increase during similar LTP induction paradigm (Meyer et al., 2014). These changes may mediate the increased presynaptic release probability during LTP (Dolphin et al., 1982; Enoki et al., 2009). Presynaptic functional potentiation likely requires retrograde signaling through messengers released by spines, such as brain-derived neurotrophic factor (BDNF) (Edelmann et al., 2015; Hedrick et al., 2016) and nitric oxide (NO) (O’Dell et al., 1991; Schuman and Madison, 1991).

Overall, our ultrastructural study revealed a timecourse of coordinated post-, extra- and pre-synaptic ultrastructural changes during LTP induced at a single spine. The combination of rapid and sustained increases in PSD complexity and eASI membrane, as well as slow development of PSD size and presynaptic function, are all likely substrates of synaptic potentiation at different temporal phases (Fig. 6).

**Figure 6.**
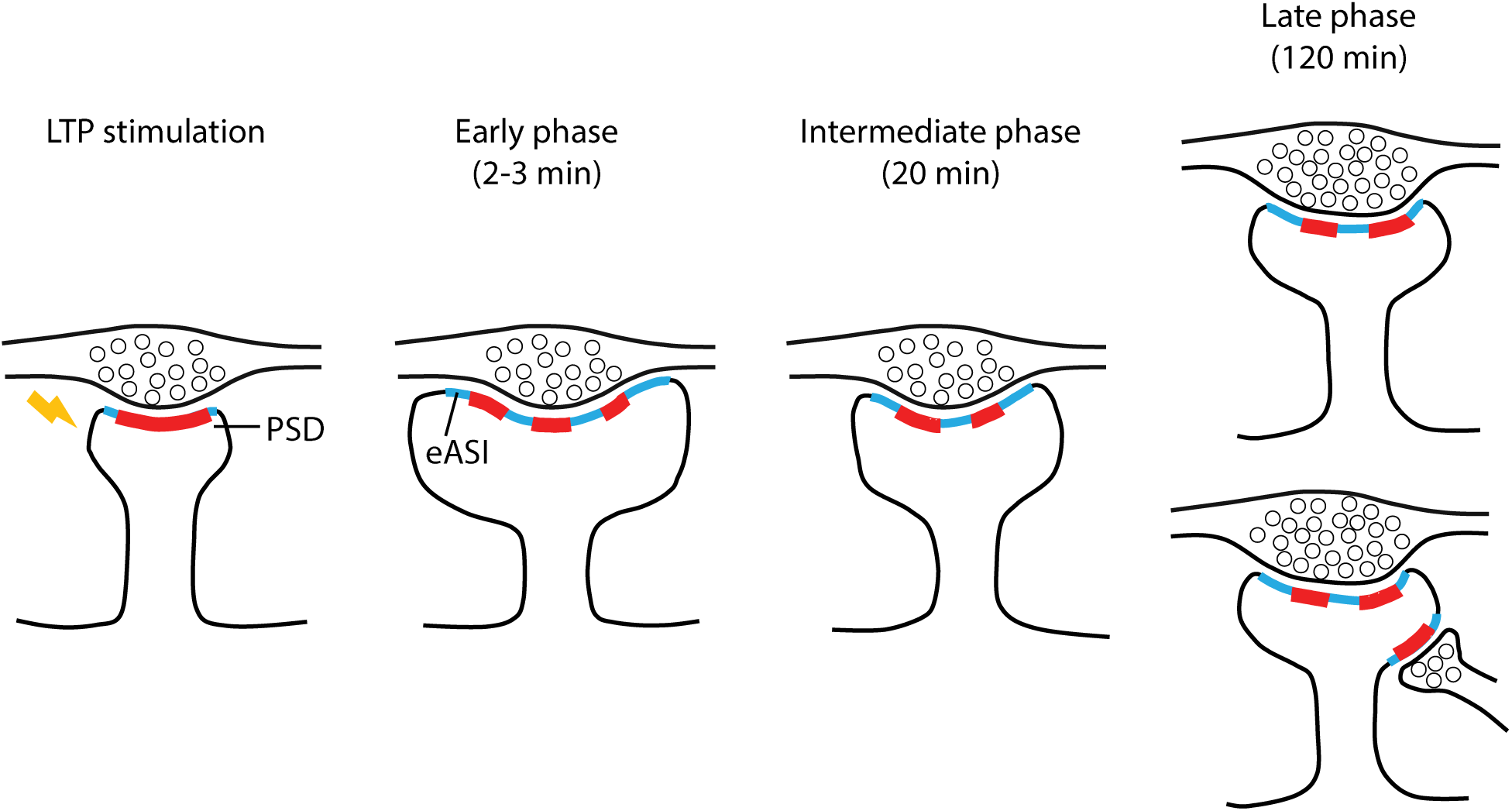
Schematic diagram illustrating the ultrastructural adaptations after LTP induced in single dendritic spines. Stimulated dendritic spines undergo dramatic spine head volume expansion and PSD (red) shape changes including segmentation and perforation immediately after LTP induction, together with the expansion of extrasynaptic axon-spine interface (eASI) area (cyan). Increased PSD complexity and eASI area maintain to the late phase of LTP. Additional presynaptic bouton might be recruited to LTP spines, forming multi-innervating spines at intermediate and late phases. Presynaptic vesicle number increases in late phase LTP, indicating enhanced presynaptic strength.

## Material and methods

### Hippocampal organotypic slice culture and gene gun transfection

All experiments were performed in accordance with the animal welfare laws of the Max Planck Florida Institute for Neuroscience Institutional Animal Care and Use Committee. Hippocampal organotypic slice cultures were prepared from P4 to P6 mice as previous described (Gogolla et al., 2006). After 10 to 12 days in culture, CAG-EGFP plasmid (purchased from Addgene, #16664) transfection was performed by a Helios gene gun (Bio-Rad, CA) (Woods and Zito, 2008). 2 days after transfection, slices with sparsely labeled CA1 pyramidal neurons were picked up for glutamate uncaging.

### 2P glutamate uncaging

MNI-glutamate uncaging was performed with 2P laser microscopy (720 nm) with 4 mW power, 0.5 Hz × 30 stimulation for LTP induction. Buffer for 2P imaging and uncaging is Mg^2+^ free ACSF containing 127 nM NaCl, 2.5 mM KCl, 25 mM NaHCO_3_, 1.25 mM NaH_2_PO_4_, 4 mM CaCl_2_, and 25 mM D-glucose plus 1 µM TTX and 2 mM MNI-glutamate aerated with 95% O_2_ and 5% CO_2_ (Patterson and Yasuda, 2010). Target spines were selected on CA1 pyramidal neuron secondary dendrites (Fig. S1A1, A2). Uncaging was performed at approximately 0.5 µm from the tip of selected spine (Fig. 1B, S1A3). 1 × (z-stack, 5 µm/step), 5 × (z-stack, 3 µm/step), and 25 × (z-stack, 1 µm/step) images of the whole neuron or specific uncaged region were taken before uncaging with 2P microscopy, 25 × images of the same region were taken immediately after uncaging to confirm the success of stimulation (Fig. 1C, S1A4). Hippocampal slices were transferred into fixative buffer at different time points (2-3 min, 20 min, 2 h) after uncaging and kept for 1h. Fixative buffer contains 2% paraformaldehyde and 2% glutaraldehyde in 0.1 M phosphate buffer (PB) at pH 7.4. Slices were then washed in 0.1 M PB 3 times for total 30 min. 25 × images of target spines were taken to further confirm the induced dendritic protrusion and maintenance of LTP (Fig. S1B1).

### Confocal scanning microscopy imaging

Confocal images were taken with a Zeiss LSM780 confocal scanning microscope (Zeiss, Oberkochen, Germany). GFP positive neurons were viewed with a laser excitation wavelength of 488 nm. Images were taken in tile mode to cover the whole slice, and in Z-stack mode (5 µm interval) to cover the Z-axis range of target neuron (Fig. S1B3).

### Laser burning marks induction

Some slices were chosen to test additional fiducial marks for correlation. For these slices, 2P laser was used to make burning marks near the target neurons and target secondary dendrites after slice fixation. For easy operation, uncaging laser at 720 nm (30 mW) was used in line scan mode to introduce horizontal burning lines while slices kept perfused in 0.1 M PB (pH 7.4). Times of line scanning need to be tested for each sample. Successful burning marks show auto-fluorescence under 2P microscopy (Fig. S1B2), and can be visualized after slice embedded in resin under both light microscopy and EM.

### Immuo-EM processing and ultra-thin sectioning

Slices were incubated in 50 mM Glycine in 0.1 M PB for 10 min to block excessive aldehyde sites. Slices were then cryo-protected with 15%-30% sucrose, followed by 2 freeze-thaw cycles of 1 min liquid nitrogen permeation. Slices were then blocked with normal goat serum and fish skin gelatin, incubated with anti-GFP primary antibody (0.1 µg/ml, ab6556, Abcam, Cambridge, United Kingdom) for 40 h and nanogold conjugated secondary antibody (1:100, #2003, Nanoprobes, NY) for 16 h. Silver enhancement was then performed with HQ silver enhancement kit (#2012, Nanoprobes, NY) to increase the visibility of gold particles. Slices were then post-fixed with 0.5% aqueous OsO4 for 40 min at 4 °C, stained with 1% aqueous uranyl acetate for 35 min, and dehydrated with gradually increased concentration of ethanol, acetone, propylene oxidase, and infiltrated by Durcupan ACE (Sigma-Aldrich, MO).

Slices were trimmed down to the region containing the target neuron according to confocal image localization and light microscopy visualization. The final sample size was around 1 mm × 1.5 mm. The samples were then sectioned with 4mm Diatome 35° knife by ATUMtome (RMC/Boeckeler Instruments, Inc., AZ) at the thickness of 50 nm, collected onto rolling Kapton tape (Fig. S1C1). The Kapton tape was then cut and aligned on carbon tape covered 4 inch silicon wafers with proper labeling. Three 150 mesh copper grids were then put onto blank regions of carbon tape to be used as fiducial markers (Fig. S1C2). A 5 nm thick layer of carbon was then coated onto the surface of wafer using a high vacuum sputter coater (ACE600, Leica, Wetzlar, Germany).

### SEM imaging

For Atlas assisted SEM imaging, an overview image of the wafer was taken with a digital camera for alignment of the physical position of the wafer in the SEM chamber (Fig. S1C2). Alignment was done by using the attached grids on the wafer as fiducial markers. Approximate locations of sections could also be viewed in the digital camera overview image. However, due to its low resolution and possible distortion, it is not accurate enough to locate and image sections with only the overview image. Then low resolution (2 µm/pixel) images of whole sections were first taken to accurately locate sections’ position by secondary electron (SE2) detector due to its high performance in surface detection and imaging speed (Fig. S1D1).

Relative location of the immunogold labeled target neuron in the serial sections was then estimated according to confocal image. A backscattered electron (BSE) detector was used at this step to search for the silver enhanced nanogold particles label profiles, and obtaine medium resolution (30 nm/pixel) images of the target neuron (Fig. S1D2). A rough 3D reconstruction of the neuron containing its soma and target secondary dendrite regions was made by Amira software (Thermo Fisher Scientific, Waltham, MA). By comparing the 3D reconstruction shape of labeled neuron and the originally captured 2P microscopy image, target neuron was finally confirmed. The 3D reconstruction was also used to determine the range of sections containing target secondary dendrite and modulated region.

Final high resolution images (4 nm/pixel) were then taken using an In-lens Duo detector in BSE mode to visualize ultrastructure of the target spines and surrounding regions (Fig. S1D3). Ultrastructure required for analyzing spines and synapses, such as PSD, presynaptic vesicle, are clearly visualized. 3D reconstruction of dendritic spines was done in Amira (Fig. S1D4).

### Image analysis

ASI was traced as the postsynaptic part of the interface between spine head and the opposing presynaptic bouton. PSD was identified as the electron-dense region in ASI, and eASI was identified as the ASI membrane without PSD.

### PSD complexity

Dendritic spines and PSDs were reconstructed in 3D in Amira, then exported to Matlab (Mathworks, MA) to create shadowed PSD region on spine surface. Smoothed PSD area and perimeter were measure and used to calculate complexity index with the following formula:

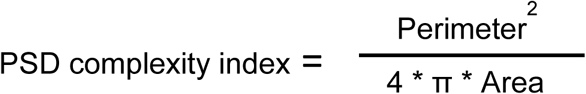

### Statistical Analysis

Statistical tests were done in Graphpad Prism 6 (Graphpad Software, CA). Measured ultrastructural data were first tested for normal distribution and lognormal distribution with D’Agostino and Person test. Data follow normal distribution were compared with Welch’s t test, and shown with mean and SEM for each control v.s. LTP group. Data follow lognormal distribution were log transformed and compared with Welch’s test. Data not follow normal or lognormal distribution were compared with Mann-Whitney test, and shown with median and interquartile range (IQR).

## Author Contributions

Y.S. designed and performed experiments, analyzed data, and wrote the paper. M.S. wrote Matlab code for data analysis. N.K. supervised EM experiments. R.Y. designed and supervised the projects, wrote Matlab code for data analysis, and wrote the paper.

## Acknowledgements

We thank Jessica Martin for performing some of the initial 3D reconstruction, Jaime Richards for preparing organotypic slice cultures, Takayasu Minuki and Debbie Guerrero-Given for technical assistance, Lesley Colgan for critical reading. We thank Max Planck Florida Institute for Neuroscience Imaging Center for instrument and technical support. This work was supported by NIH (DP1NS096787, R01MH080047).

---

**Figure S1.**
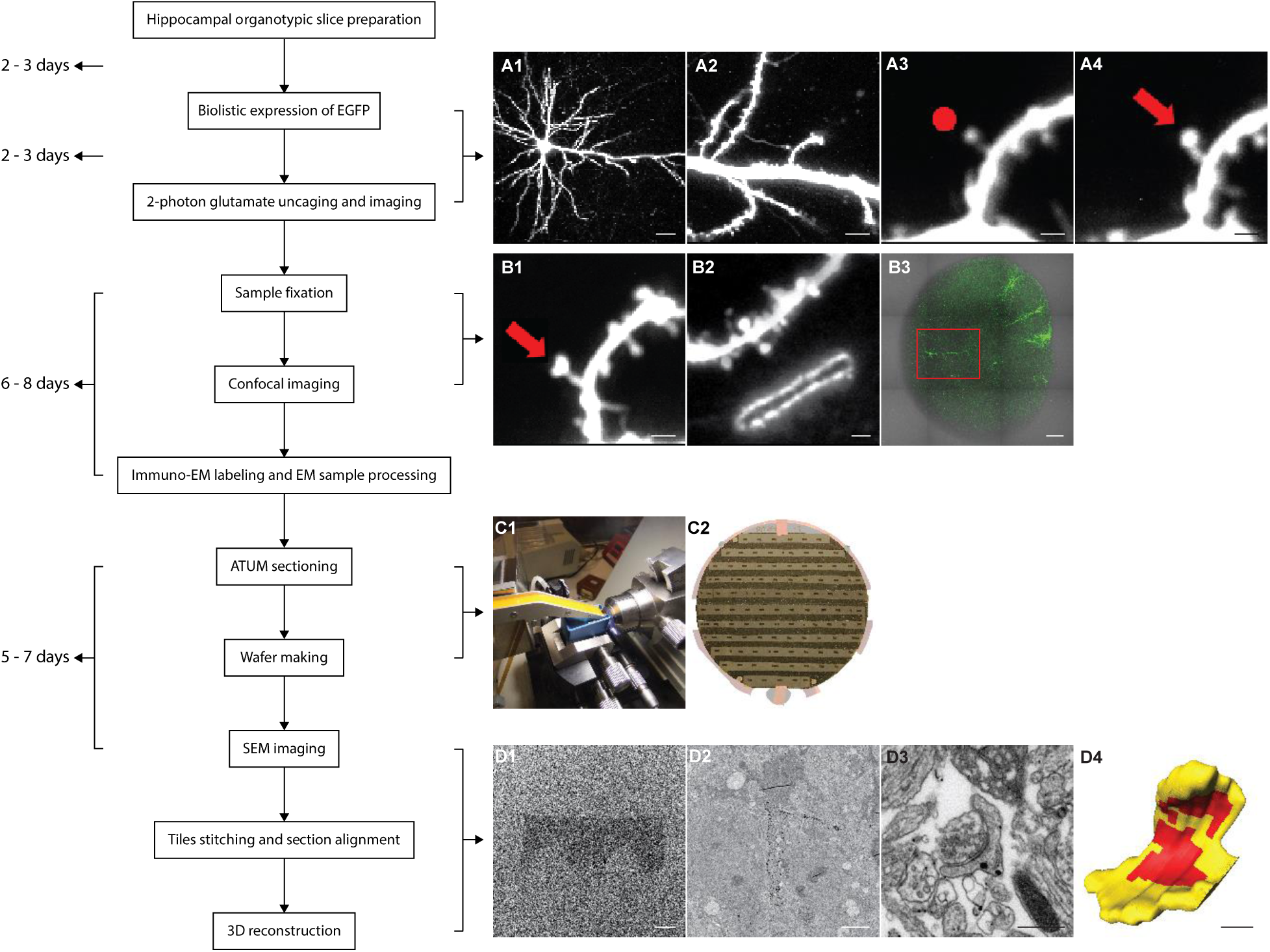
Correlative light and electron microscopy imaging of a 2P uncaging modulated dendritic spine. **(Left panel)** Workflow and estimated cost time **(A1-A2)** 1 x and 5 x 2P images of a GFP positive CA1 pyramidal neuron in organotypic hippocampal slice selected for glutamate uncaging. Scale bar: A1, 20 µm, A2, 5 µm. **(A3-A4)** 25 x 2P images of a dendritic spine before **(A3)** and after **(A4)** 2P glutamate uncaging stimulation. Red dot in **(A3)** shows the uncaging position, and red arrow in **(A4)** points to the enlarged dendritic spine. Scale bar: 1 µm. **(B1)** 25x 2p image of the uncaged spine after fixation. Arrow points at the target spine. Scale bar: 1 µm. **(B2)** Laser burning fiducial mark is introduced by 2p laser near the target spine after tissue fixation. Scale bar: 1 µm. **(B3)** A confocal image shows the overview of GFP expression pattern in the whole hippocampal organotypic slice. Target neuron is labeled in red square. Scale bar: 200 µm. **(C1)** ATUMtome setup for sectioning and collecting ultrathin sections. **(C2)** Silicon wafer holding aligned Kapton tape containing serial sections. 3 copper grids on the edge of the wafer are used as fiducial markers for alignment. **(D1)** 2 µm/pixel image taken with SE2 detector shows the outline of two ultrathin sections. Scale bar: 200 µm. **(D2)** 30 nm/pixel image taken with BSE detector shows tissue structure and the silver enhanced nanogold labeling pattern. Nanogolds are seen as black dots in the image. Scale bar: 50 µm. **(D3)** 4 nm/pixel high resolution image taken with InlensDuo detector shows details of modulated dendritic spine ultrastructure. Scale bar: 0.5 µm. **(D4)** 3D reconstruction of the target dendritic spine. Spine head colored in yellow, and PSD colored in red. Scale bar: 200 nm.

**Figure S2.**
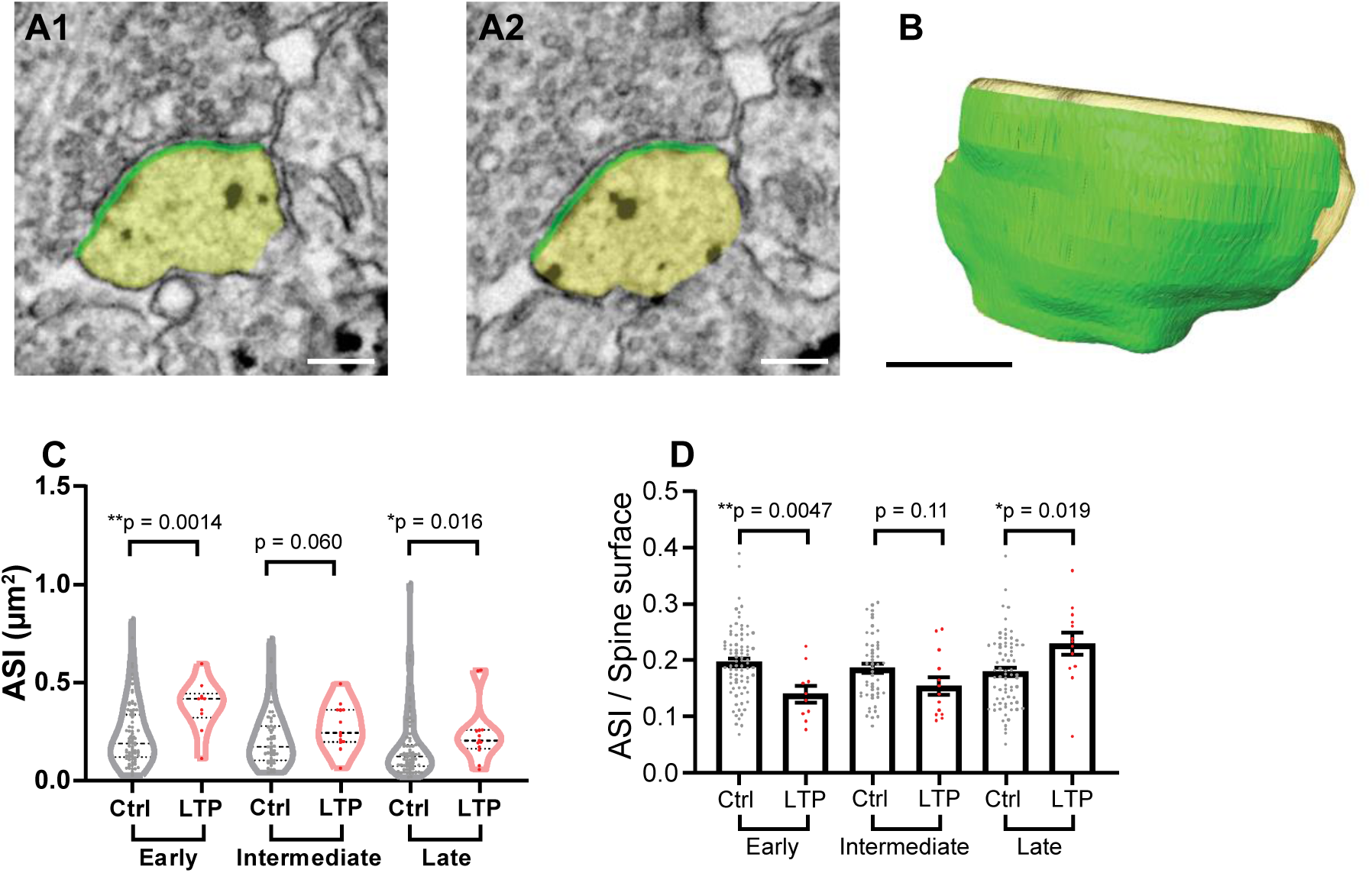
LTP induces expansion of Total ASI postsynaptic membrane. **(A-B)** Serial images of an example spine **(A)** and 3D reconstruction of the example spine **(B)** with spine head colored in yellow, and ASI colored in green. Scale bar: 200 nm. **(C)** ASI area size at early, intermediate and late phases. Welch’s t-test on log-transformed data. **(D)** Ratio of ASI area to spine surface at early, intermediate and late phases. Welch’s t-test, mean ± SEM shown.

